# Remodeling gut microbiota by *Streptococcus thermophilus* 19 attenuates inflammation in septic mice

**DOI:** 10.1101/543967

**Authors:** Fu Han, Yijie Zhang, Xuekang Yang, Zhuoqun Fang, Shichao Han, Xiaoqiang Li, Weixia Cai, Dan Xiao, Jiaqi Liu, Wanfu Zhang, Gaofeng Wu, Dahai Hu

**Author notes:** Address correspondence to Gaofeng Wu, and Dahai Hu,. Fu Han, Yijie Zhang and Xuekang Yang contributed equally to this work.

## Abstract

Sepsis is a life-threatening organ dysfunction caused by a dysregulated host response to infection and is the leading cause of death in burn patients. *Streptococcus thermophilus* 19 is a highly effective probiotic, with well-studied health benefits, but its role in protecting viscera against injury caused by sepsis and the underlying mechanism is poorly understood. The goal of this study was to evaluate protection potency of S. *thermophilus* against inflammation in mice and evaluate the influence of sepsis and S. *thermophilus* on microbial community. We tested the utility of S. *thermophilus* 19 in attenuating inflammation in *vitro* and *vivo* of LPS-induced sepsis mouse model. We also evaluated the influence of sepsis and S. *thermophilus* on microbial community. In *vitro,* S. *thermophilus* 19 decrease the expression of inflammatory factors. Additionally, in a lipopolysaccharide-induced septic mouse model, mice administered the probiotic 19 was highly resistant to Lps and exhibited decreased expression of inflammatory factors compared to Lps-treated control mice. A MiSeq-based sequence analysis revealed that gut microbiota alterations in mice intraperitoneally injected with 1 mg/ml LPS were mitigated by the administration of oral probiotics 19. Together these findings indicate that S. *thermophilus* 19 may be a new avenue for interventions against inflammation caused by sepsis and other systemic inflammatory diseases. In an analysis of the gut microbiota of the all group mice, we found that sepsis is associated with gut microbiota and probiotics attenuate the inflammation through remodeling gut microbiota.

**Importance:** Sepsis is life-threatening organ dysfunction which is the leading cause of death in burn patients. Although our understanding of sepsis has increased substantially in recent years, it’s still reported to be the leading cause of death in seriously ill patients. Evidences showed that gut microbiota play an important role in sepsis. Moreover, probiotics have been used to prevent numbers of gut health disorders and alleviate inflammation associated with some human diseases by promoting changes in the gut microbiota composition. Hence, to investigate the mechanism of probiotics in the treatment of sepsis has emerged. The significance of our research is in identifying the role of gut microbiota in sepsis and found an effective probiotic that reduces inflammation, S. *thermophilus* 19, and investigating the therapeutic effect and mechanism of S. *thermophilus* 19 on sepsis, which might be a new avenue for interventions against inflammation caused by sepsis and other systemic inflammatory diseases.

## Introduction

Sepsis is a life-threatening organ dysfunction caused by a dysregulated host response to infection and is the leading cause of death in burn patients, responsible for up to 50 to 60% of burn injury deaths (1, 2). Although our understanding of sepsis has increased substantially in recent years, it is still reported to be the leading cause of death in seriously ill patients, and the incidence of sepsis has increased annually. Therefore, new insights into the causes of sepsis are urgently needed.

The gut microbiota is a complex ecosystem consisting of trillions of bacteria that live in the digestive tracts of humans and other animals (3). Growing evidence supports the key role of a healthy gut microbiota in promoting and maintaining a balanced immune response and in the establishment of the gut barrier immediately after birth (4, 5). Moreover, a dysbiotic state of the gut microbiota can lead to dysregulation of various processes, which can in turn contribute to the development of autoimmune conditions (6). For instance, the presence or overabundance of specific types of bacteria may contribute to inflammatory disorders such as IBD (6). Additionally, metabolites from certain members of the gut flora may influence host signaling pathways, contributing to disorders such as colon cancer and obesity. Sepsis is an extreme response to inflammation that has profound effects on all parts of the body. For decades, the gut has been regarded as the motor of sepsis (7), and it has recently been shown that a healthy gut microbiota has a protective role during systemic inflammation. Thus, we hypothesized that intestinal bacteria play an important role in sepsis since the gut microbiota is associated with many diseases.

Probiotics are live microbes that have beneficial effects on human and animal health when ingested in sufficient amounts (8). Probiotics play an important role in maintaining the normal microbiota composition and have been used to treat or prevent a number of gut health disorders, such as irritable bowel syndrome, hypercholesterolemia, gastritis, gut infection, parasitic infestation, hypersensitivity (including food allergies), and even certain types of cancers (e.g., colorectal cancer) (9, 10). The use of microbes as probiotics also hold potential for oral health in preventing and treating oral infections, dental plaque-related diseases, periodontal diseases and halitosis. Furthermore, probiotics can alleviate inflammation associated with some human diseases by promoting changes in the gut microbiota composition (11, 12). *Streptococcus thermophilus* is a highly effective probiotic that has well studied health benefits, including the production of antibiotics that prevent infections from pneumonia-causing microbes and C. *difficile* and can help to prevent ulcers (13–15).

In this study, we used a coculture system (probiotics and RAW264.7 cells) to assess the ability of probiotics to decrease the expression of inflammatory factors. We showed that *Streptococcus thermophilus* 19 can decrease the inflammation induced by Lps in RAW264.7 cells. Furthermore, we investigated the ability of S. *thermophilus* 19 to protect mice against Lps-induced inflammation and gut microbiota alterations when administered as a probiotic. We observed that the administration of S. *thermophilus* 19 as probiotics could alter the gut microbiota composition of untreated mice or mice with Lps-induced sepsis, with the symptoms of sepsis mitigated in the latter group. Moreover, the levels of several inflammatory factors in various organs were correlated to a diverse gut microbiota composition. We hypothesize that supplementation of diets with probiotics protects visceral organs by reducing inflammation through alterations in the gut microbiota after sepsis.

## Results

### Probiotics decrease the expression of inflammatory factors in vitro

To assess the influence of the assayed probiotics on the expression of inflammatory factors, we developed a co-culture system (probiotics and RAW264.7 cells). After incubating for 6 hours, total RNA was extracted and the expression of inflammatory factors was assessed via quantitative RT-PCR. The Lps treatment increased the expression of inflammatory factors compared to the untreated group. After co-culturing with probiotics, we observed a reduction in inflammatory factor expression, particularly when cells were incubated S. *thermophilus* 19 (Figure 1). At the same time, we investigated the influence of S. *thermophilus* 19 on the cell viability after treatment 6 hours. Results showed that S. *thermophilus* 19 didn’t affect the cell viability after co-culture 6 hours (Supplementary Figure1). Therefore, S. *thermophilus* 19 was chosen for further study.

**Figure. 1.**
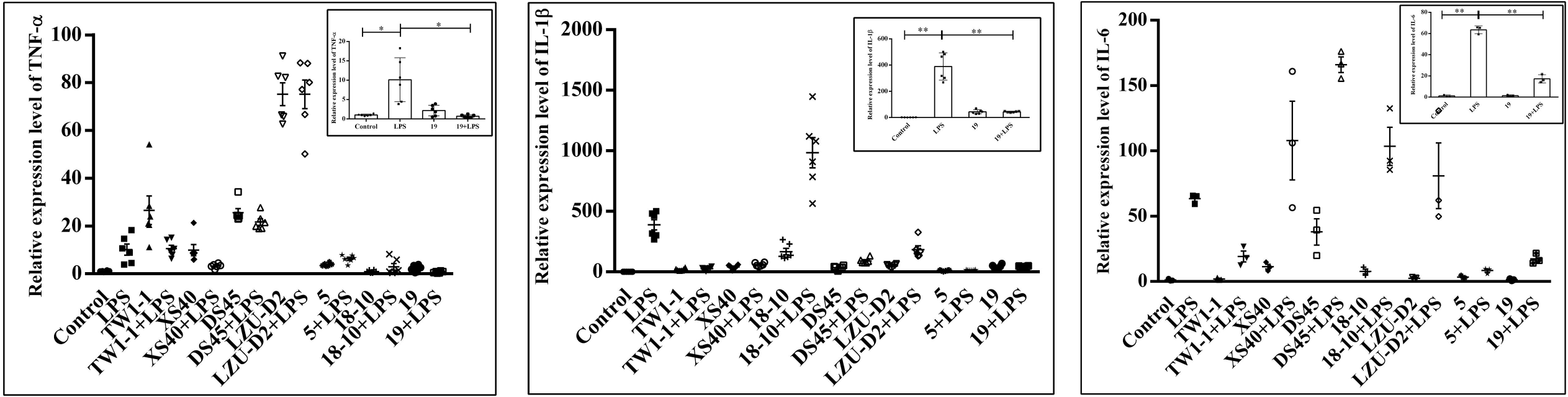
The expression of inflammatory factors (IL-1β, TNF-α and IL-6) in Lps-treated, probiotics-Lps treated and untreated RAW264.7 cells. Error bars represents SEM. Illustration represent the influence of S. *thermophilus* 19 on inflammatory factors. *P<0.05, **P<0.01.

### Probiotics effectively alleviated inflammation induced by sepsis

At first, the influence of different doses of Lps on mice survival rate was investigated. All mice died when the concentration of Lps exceeded 2.5 mg/kg, even in mice administered probiotics (data not shown). However, nearly 60% of mice survived when administered probiotics together with 2 mg/ml Lps, whereas only 20% of mice administered the same Lps without probiotics survived (Figure 2A). All mice treated with 1 mg/kg of Lps survived (Figure 2A). So, 1mg/kg Lps was chosen to investigate the influence of S. *thermophilus* 19 on gut microbiota and inflammation of sepsis. Mice treated with Lps lost approximately 10% of their body weight during the 48 hours after injection, while untreated mice did not lose weight (Supplementary Figure2A). Although all treated groups regained their baseline weight by the third day, mice treated with S. *thermophilus* 19 exhibited high rates of body weight recovery. Total food and water intake and the animal health conditions for all mice were recorded. The reason we recorded the total water and food is we keep one group of mice in a cage. Lps-treated mice with or without probiotics exhibited a reduction in total drinking water and rat chow intake (Supplementary Figure2B). Furthermore, mice treated with the probiotics alone also exhibited decreased water and rat chow intake (Supplementary Figure 2B). However, mice treated with S. *thermophilus* 19 alone showed no changes in body weight, although they exhibited lower drinking water and food intake (Supplementary Figure2A and 2B). The decrease in body weight of the Lps-treated mice could be explained by the Lps-induced inflammation causing a reduction in food and drinking water intake, while the probiotics could alleviate inflammation to promote the recovery in body weight.

**Figure. 2.**
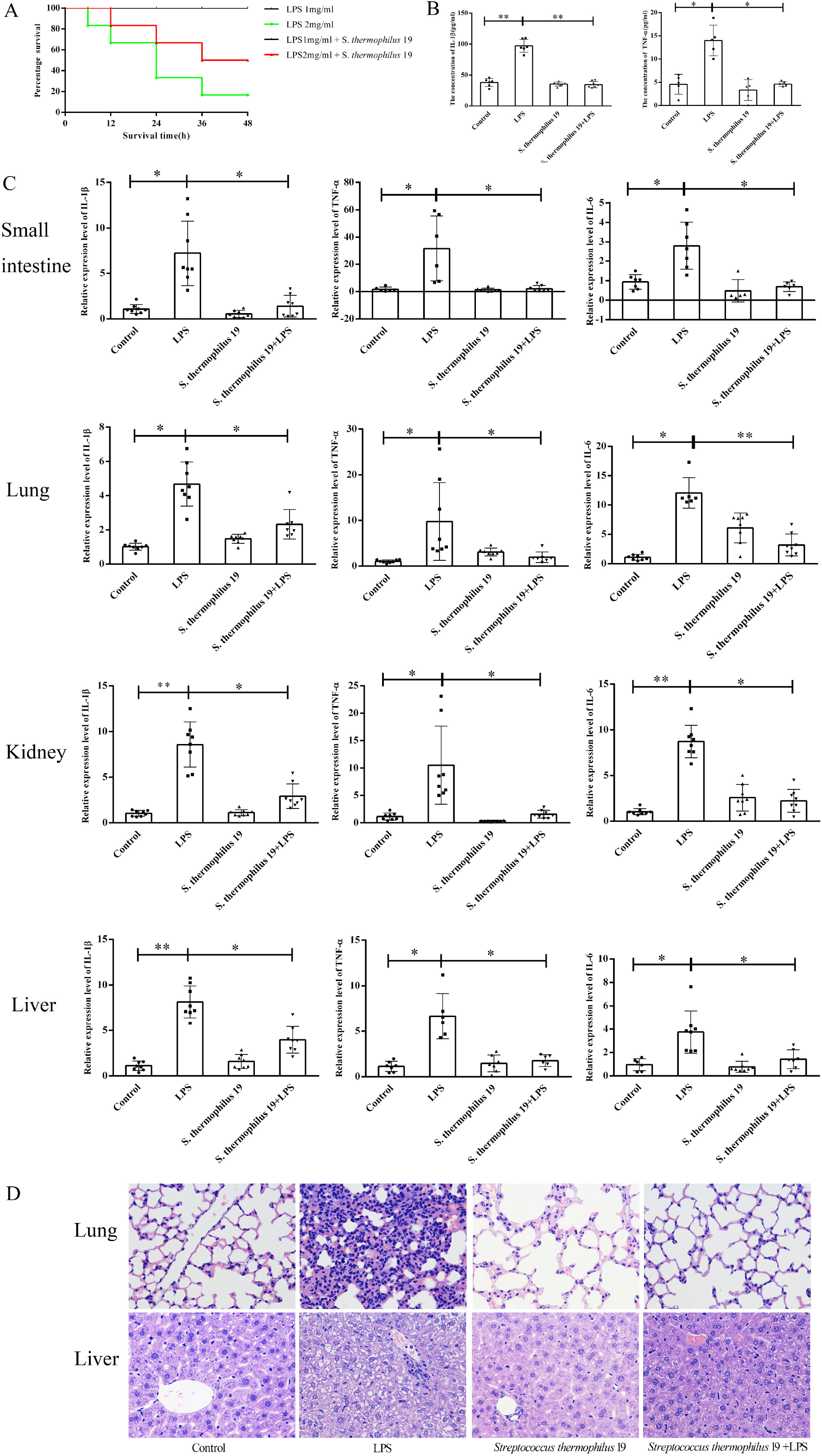
Probiotics alleviate the inflammatory caused by Lps-induced sepsis. (A) Survival rates of mice with or without probiotics treatment after 48h stimulation with different dose of Lps (n=10). (B) Levels of IL-1β and TNF-α in blood were determined using commercial ELISA kits (n=8). (C) Probiotics intervention resulted in decreased inflammation small intestine, lung, liver and kidney (n=8). Error bars represents SEM. (D) Hematoxylin and eosin staining of liver, and lung tissues from different groups. Sections were examined and photographed under a microscope.

We observed a 2-fold increase in TNF-α expression and 2.5-fold in IL-1β while it was reduced to that observed in the control group in mice administered probiotics (Figure 2B). In contrast, in mice treated with probiotics without Lps treatment, no significant effect on the serum levels of IL-1β and TNF-α were observed compared to the control group, demonstrating that the probiotics has no influence on the host in the absence of sepsis.

Next, inflammation state of the kidneys, small intestines, livers and lungs of each mouse after Lps and probiotic treatment was investigated. Lps treatment dramatically increased the expression of IL-1β, IL-6 and TNF-α in all tissues while they were effectively rescued in the mice treated with S. *thermophilus* 19 compared to the mice treated with Lps alone (Figure 2C). However, the expression of TNF-α, IL-6 and IL-1β in the probiotic- -treated mice and control mice did not significantly differ (Figure 2C). H&E staining revealed that compared with the liver sections in control group mice, significant congestion of veins and hepatocyte necrosis was observed in the Lps-treated mice, and the loss of intact liver plates and hepatocyte vacuolization was observed (Figure 2D). In pulmonary sections, drastic destruction of alveolar structures was detected in the Lps-treated mice, and the effusion in alveoli in these mice was markedly more severe than that observed in the control group mice. Furthermore, tissue infiltration by inflammatory cells was substantially higher in Lps-treated mice than in the control group mice. Co-treatment with probiotics resulted in the restoration of a close-to-normal appearance of liver and lung tissues. Moreover, S. *thermophilus* 19 treatment alone did not affect the liver and lung sections of mice (Figure 2D).

### Lps altered the gut microbiota structure of mice

In the balance between gut microbiota and inflammation, deviations either way may cause corresponding adjustments in the other. To test whether the gut microbiota of mice was altered due to sepsis, we collected cecal feces of mice and assayed them via MiSeq sequencing to determine the composition of gut microbiota. Mice treated with Lps exhibited decreases gut microbiota richness compared to the control group (Chao1 index) (P<0.05) while no difference in diversity (Shannon index) between two groups was observed (Figure 3A and 3B). Gut microbiota of mice treated with Lps only clustered differently from those of the control group mice, demonstrating the significant effect of Lps on the gut microbiota (Figure 3C). The relative abundance of gut microbiota in control and Lps group was showed in Figure 3D. In details, compared to the control group, Lps-treated mice had lower abundances of bacteria belonging to the phylum *Fusobacteria* and of the genera *Fusobacterium* and *Psychrobacter* (Supplementary Figure3A and 3B) (P<0.05). In contrast, higher abundances of bacteria from the genus *Flavonifractor* were observed in the Lps-treated mice. Interestingly, 8 OTUs were specifically present in the Lps-treatment group compared to the control group, while the control group also contained 8 specific OTUs (Supplementary Figure 4).

**Figure. 3.**
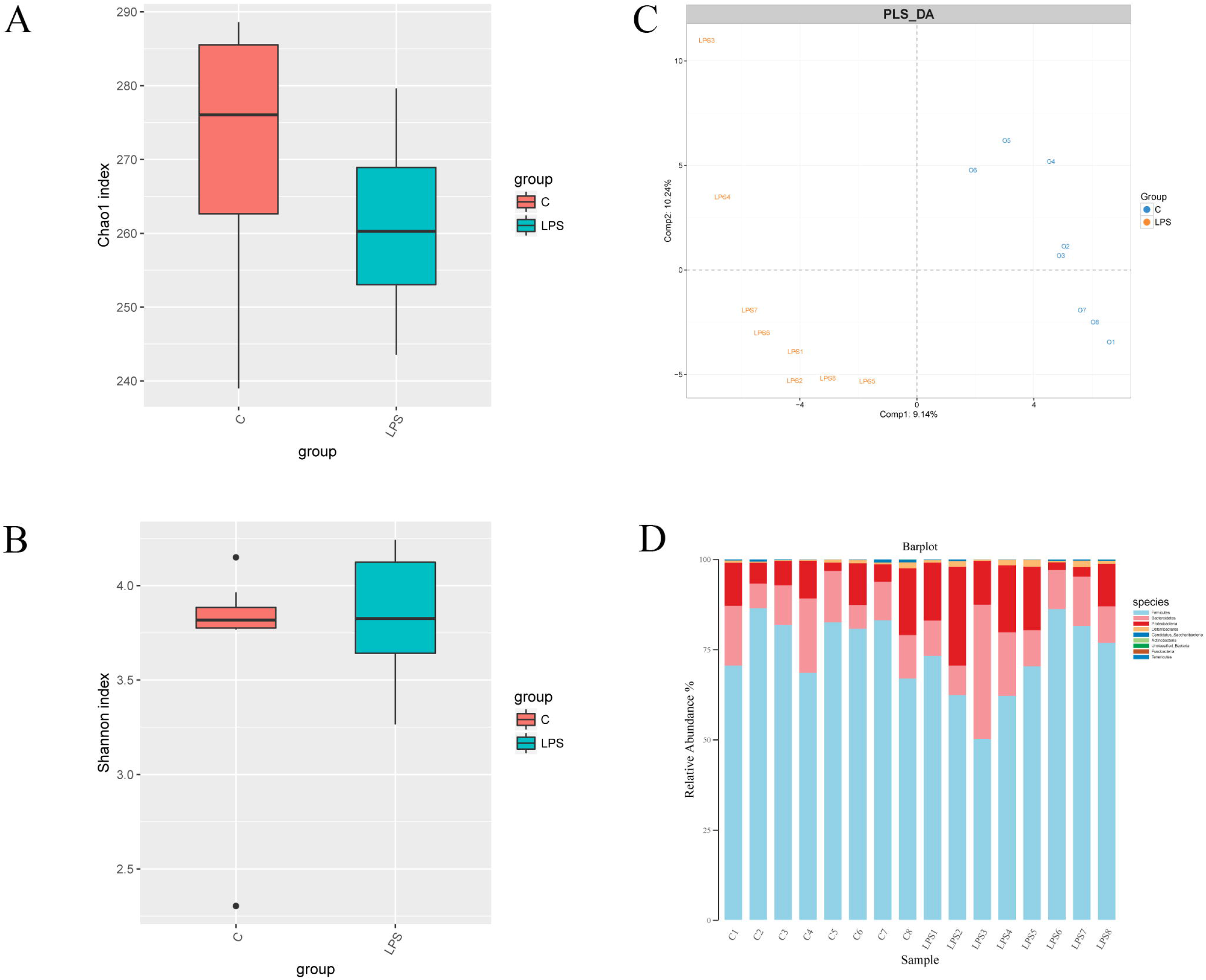
Lps induce significant impact on microbiota composition. (A) (B) Fecal microbiota alfa diversity. (C) PLS_DA plot of fecal microbiota of Lps-treated or control mice. (D) The change of gut microbiota at phylum level.

### Probiotics intervention alters the gut microbiota of mice

To investigate the effect of probiotics on the gut microbiota of mice, we sequenced the gut microbiota of the mice treated with probiotics alone. The diversity of gut microbiota differed for the various probiotics assayed compared to control group (P<0.05), while no difference in richness was observed among the groups (Figure 4A and 4B). Moreover, gut microbiota of mice treated with probiotics alone clustered differently from that of control group mice (Figure 4C). The relative abundance of gut microbiota in control and Lps group was showed in Figure 4D. In details, mice treated with S. *thermophilus* 19 exhibited a decreased abundance of bacteria belonging to the phylum *Bacteroidetes* and an increased abundance of the phylum *Firmicutes.* The changes in microbiota compositions in the 19 treatment mice is shown in Supplementary Figure 3A and 3B (P<0.05). Nine OTUs were specifically present in the group treated with S. *thermophilus* 19 alone compared to control group, while control group also had 5 specific OTUs (Supplementary Figure 4).

**Figure. 4.**
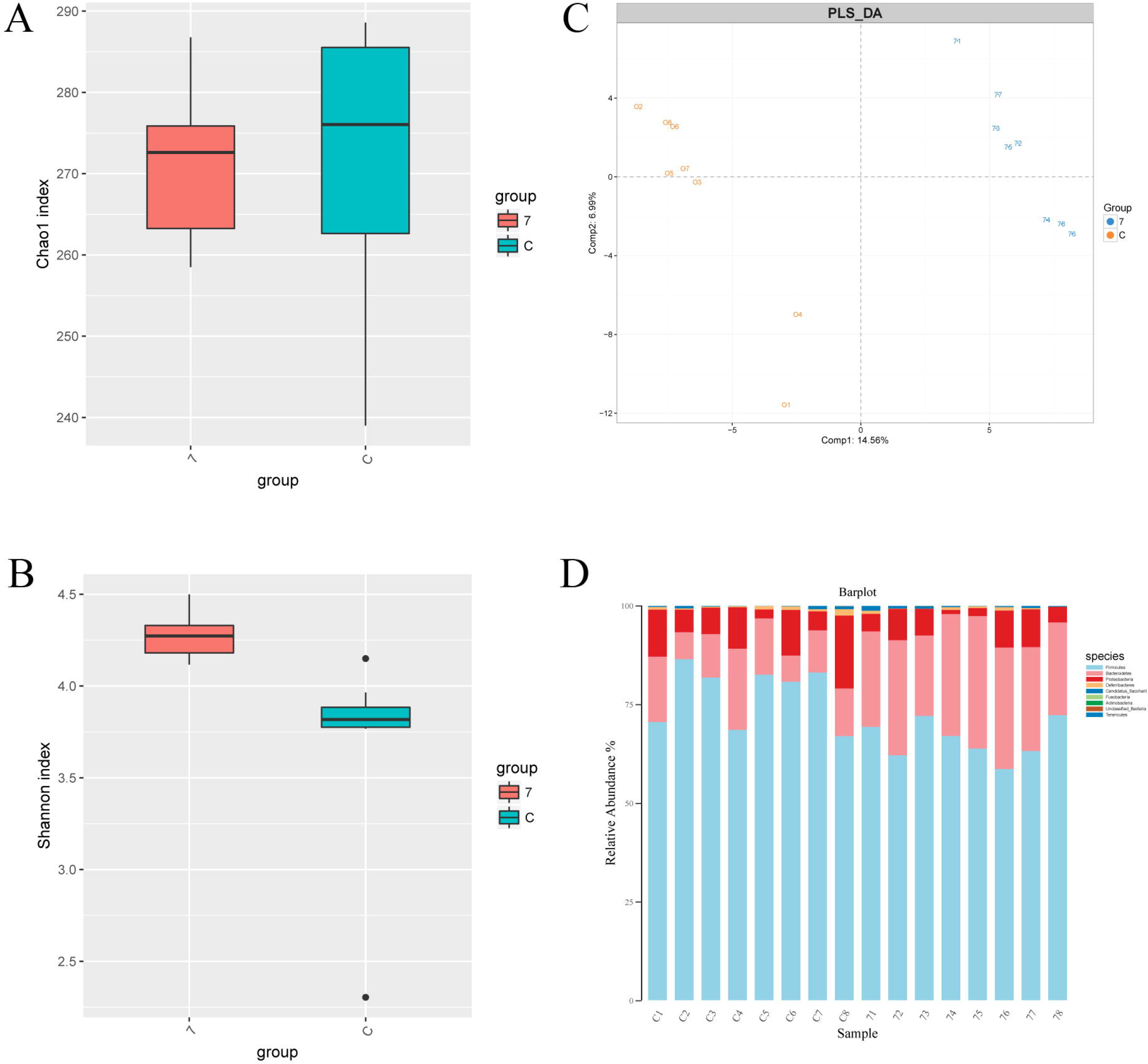
S. *thermophilus* 19 induce significant impact on microbiota composition compared to control group mice. 7 represent S. *thermophilus* 19 (n=8). (A) (B) Fecal microbiota alfa diversity. (C) PLS_DA plot of fecal microbiota of LPS-treated mice with or without S. *thermophilus* 19 treatment. (D) The change of gut microbiota at phylum level.

### Oral administration of Probiotics alleviated viscera damage via altering the gut microbiota

We showed that probiotic intervention can attenuate the inflammation in septic mice (Figure 2). Furthermore, we previously reported that probiotics can reduce the inflammation induced by Cr (VI) in mice through modifying the gut microbiota. Thus, we hypothesized that the protection of viscera by the probiotic-induced attenuation of inflammation in septic mice is also associated with changes in the intestinal microbiota. To test this hypothesis, we sequenced the 16S rRNA gene variable (V) V3-V4 region of the fecal bacteria samples obtained from S. *thermophilus* 19 treated Lps-treated mice (Lps7) and compared the results to those obtained from the mice treated with Lps alone and the control group. Overall, differences between the S. *thermophilus* 19- and Lps-treated mice were observed (Figure 5A and 5B). Meanwhile, gut microbiota of Lps+ S. *thermophilus* 19 groups clustered differently from mice treated with Lps alone group (Figure. 5C), demonstrating the important effect of probiotics. Lps-treated mice administered S. *thermophilus* 19 had lower abundance of *Clostridium*_XIVb and a higher abundance of *Fusobacterium* and *Klebsiella.* Compared to the Lps group, Lps-treated mice administered S. *thermophilus* 19 exhibited an increased abundance of *Fusobacteria* (Figure. 5D, Supplementary Figure 3A and 3B) (P<0.05). Mice administered S. *thermophilus* 19 and treated with Lps had 8 specific OTUs compared to mice treated with Lps alone (Supplementary Figure 4).

**Figure. 5.**
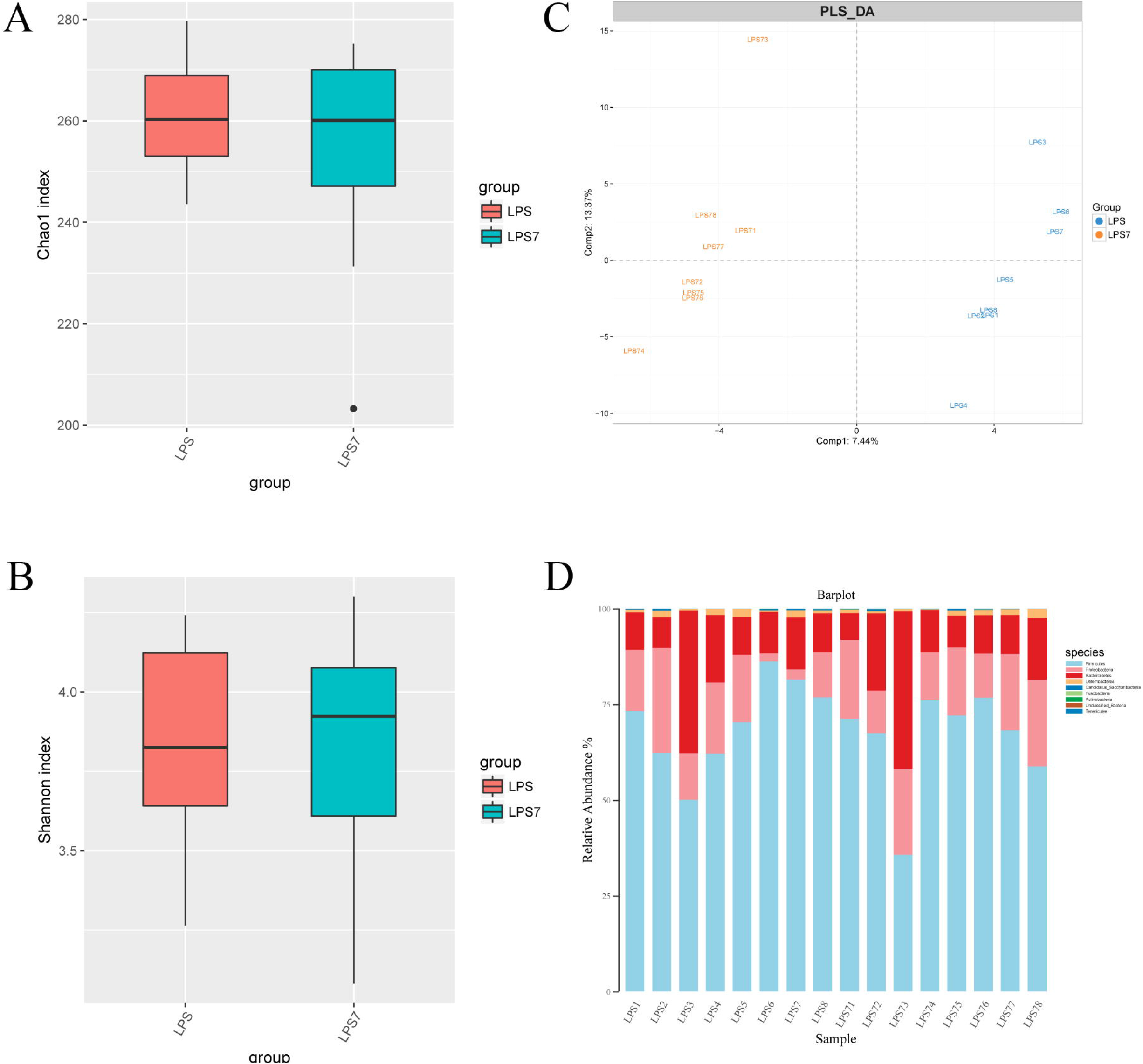
S. *thermophilus* 19 induce significant impact on microbiota composition compared to Lps-treated mice. 7 represent S. *thermophilus* 19 (n=8). (A) (B) Fecal microbiota alfa diversity. (C) PLS_DA plot of fecal microbiota of LPS-treated mice with or without S. *thermophilus* 19 treatment. (D) The change of gut microbiota at phylum level.

Next, we compared the differences in gut microbiota composition between the probiotic- and Lps-treated mice and the control group mice. Mice treated with Lps and S. *thermophilus* 19 exhibited decreases gut microbiota richness compared to the control group (Chao1 index) (P<0.05) while no difference in diversity (Shannon index) between two groups was observed (Supplementary Figure 5A and 5B). The gut microbiota of mice treated with Lps and 19 clustered differently from that of the control group mice (Supplementary Figure 5C). The change in the microbiota composition between Lps+ S. thermophilus 19 and control groups is shown in Supplementary Figure 3 (in details) (P<0.05) and Supplementary Figure 5D. Six specific OTUs were identified in the LPS7 group mice and 9 were identified in the control group (Supplementary Figure 4). Taken together, these results indicated that all the treatments altered the composition of gut microbiota of the assayed mice. Although the composition of gut bacteria in mice treated with probiotic and that of control group differed, the expression of inflammation-associated factors in these mice did not significantly differ. We speculated that the gut microbiota in these exhibited a healthy status, whereas the probiotic and Lps-treated mice had a lower health status.

Overall, these data showed that Lps and probiotics significantly impacted the microbiota composition of mice.

### The function of gut microbiota was specifically altered after the administration oral probiotics

Next, we used a Kruskal-Wallis/Wilcoxon rank-sum test to determine how the altered community structure of the gut microbiota affects its function. Mice treated with Lps and S. *thermophilus* 19 were decreased in both primary bile acid biosynthesis and secondary bile acid biosynthesis, which have proinflammatory properties compared to the Lps-treated mice (Figure 6). These data suggest a significantly decreased proinflammatory signature, as well as an increased anti-inflammatory capacity of the gut microbiome in probiotic-treated mice. Taken together, the probiotics were observed to reshape the gut microbiota with a distinct composition, network topology and functionality.

**Figure. 6.**
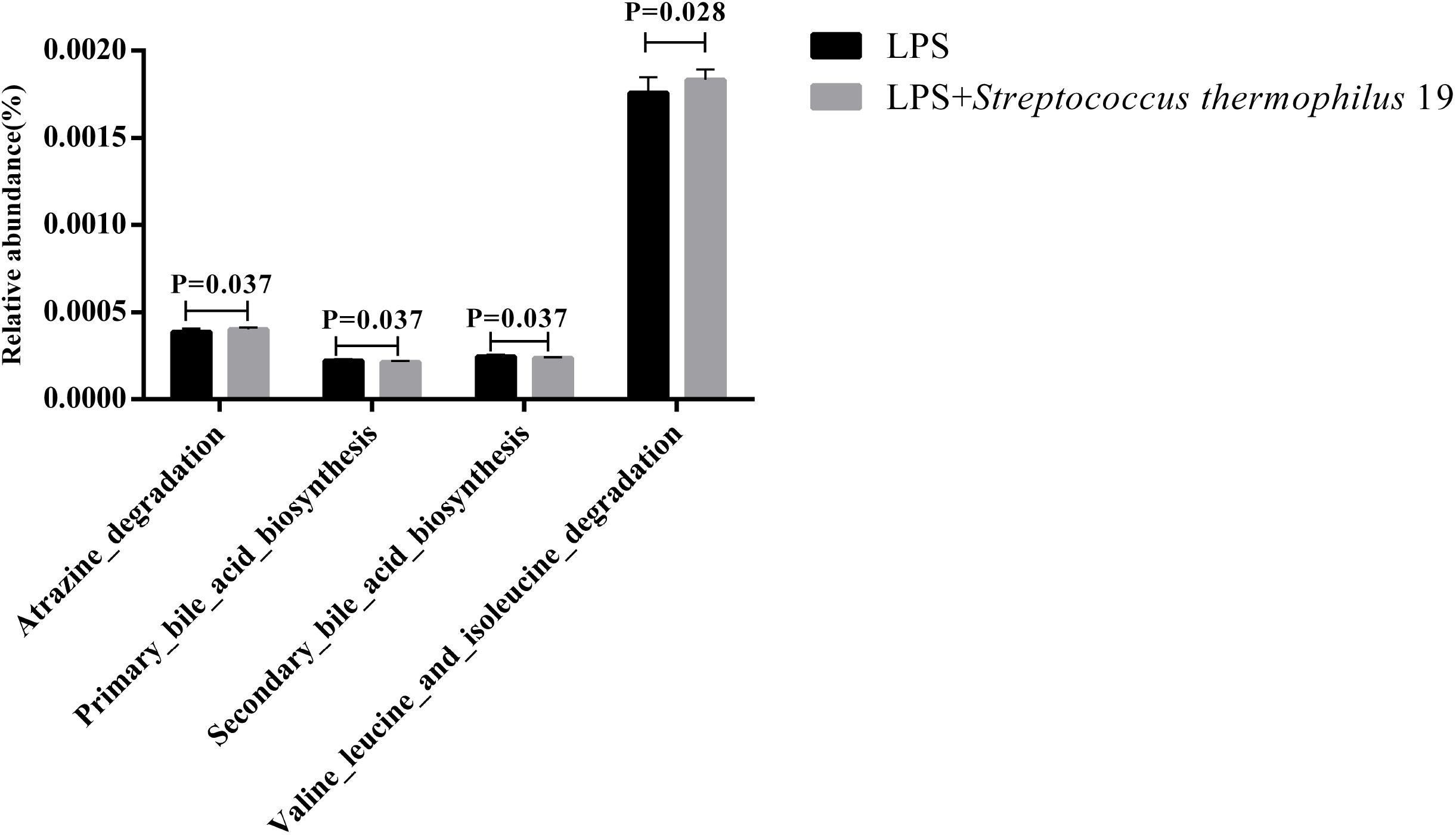
The presence of S. *thermophilus* 19 induces changes in gut microbiota function after Lps treatment. Statistical comparison was performed by first testing normality using Kruskal-Wallis/Wilcoxon rank-sum test. Error bars represents SEM.

## Discussion

Sepsis is life-threatening organ dysfunction caused by a dysregulated host response to infection and often causes multiple organ damage. S. *thermophilus* has been shown to be highly effective probiotic strains with well-studied health benefits. However, the impact of S. *thermophilus* on the gut microbiota composition, and its influence on the inflammation caused by Lps-induced sepsis remains poorly understood. In this study, we utilized a MiSeq sequencing approach to assess how S. *thermophilus* 19 modulate the host fecal microbiota and inflammatory response in an Lps-induced mouse sepsis model. Our results showed that S. *thermophilus* 19 can decrease the expression of inflammatory factors RAW264.7 cells treated with Lps. Moreover, we showed that S. *thermophilus* 19 were able to protect viscera against damage induced by sepsis. Furthermore, S. *thermophilus* 19 could alter the microbiota composition and restore homeostasis of the gut microbiota disrupted by sepsis.

Inflammation and infection are frequently accompanied by an imbalance in the intestinal microflora(16). A strong inflammatory response may then be mounted against microfloral bacteria, leading to a perpetuation of the inflammation and gut barrier dysfunction(17). Sepsis is life-threatening organ dysfunction caused by a dysregulated host response to infection, which is often causes a systemic inflammatory response. To assess the relationship between the gut microbiota and sepsis, we induced sepsis in mice through intraperitoneal injection of Lps (2 mg/ml) and used a MiSeq sequencing-based approach to evaluate the gut microbiota compositions of the assayed mice. The results showed that the Lps treatment decreased the abundance of *Fusobacteria* and the richness of the intestinal microbiota. Moreover, the abundances of the genera *Fusobacterium, Flavonifractor* and *Psychrobacter* were altered in the septic mice. Previous studies showed that shifts in the intestinal *Firmicutes* to *Bacteroidetes* ratio, as well as reduced microbiota diversity(18, 19). However, these studies had many uncertainties with regard to the variability and temporal nature of sepsis-induced dysbiosis. Thus, we used an Lps-induced sepsis model to investigate the changes in gut microbiota composition to eliminate the influence of other factors. Our results suggest that the genera *Fusobacterium, Flavonifractor* and *Psychrobacter* may play important role in the development of sepsis.

We observed that Lps significantly upregulates the expression of genes involved in inflammation, especially in the livers, lungs, kidneys and small intestines of mice. Moreover, Lps induced sepsis has been demonstrated to result in the expression of inflammation-related genes in multiple organs(1). Probiotics are live microbial food supplements or bacterial components that have been shown to have beneficial effects on human health. Additionally, probiotics are often used to treat inflammation-related diseases, such as inflammatory bowel disease, allergic diseases, and acute gastroenteritis. S. *thermophilus* is probiotics that have been used to treat many illnesses. Probiotics containing S. *thermophilus* KB19 significantly increased betaine plasma levels in chronic kidney disease(20–23). Similarly, we observed that S. *thermophilus* decreased the level of inflammatory factors in an LPS-induced sepsis mouse model. In addition, the administration of S. *thermophilus* 19 did not trigger any inflammation or dysbiosis of the gut microbiota, suggesting that they could safely be used to treat sepsis with no obvious harmful side effects. Thus, together with previous results, these results suggest that S. *thermophilus* 19 may be one alternative probiotics for use in sepsis intervention in the future.

It has now been recognized that alterations in gut microbiota composition and function appear to be an important mechanism by which probiotics alleviate human disease. Our results showed that the probiotics 19 altered the function of the gut microbiota in mice. In particular, mice treated with LPS and probiotics exhibited changes in the function of oxidative-phosphorylation and bile acid biosynthesis, which are important in inflammation-related diseases(24, 25). Moreover, probiotics also caused other functions of the gut microbiota to change. Meanwhile, mice treated with probiotics alone also exhibited changes in the function of the gut microbiota that may be good for host health by promoting low inflammatory factor expression and a good health state. Taken together, our results indicated that probiotics are good for host health despite the changes they induce in the composition and function of the gut microbiota.

In summary, we demonstrated that the probiotics S. *thermophilus* 19 can alleviate inflammation both *in vivo* and *in vitro.* This probiotics reduced the levels of inflammatory factors caused by sepsis, which may occur through multiple targets. For instance, probiotics can resistant some pathogenic bacteria enriched in gut after intraperitoneal injection of LPS, alter the functional potential of intestinal microbes, promote higher intestinal permeability, and alter the composition of the gut microbiota. These results suggest that the probiotics S. *thermophilus* 19 may be used to treat to not only sepsis but also other systemic inflammatory diseases (inflammatory bowel disease, systemic inflammatory arthritis, multiple sclerosis and so on). Collectively, the results of our study provide a conceptual framework to further text this hypothesis in humans to treat sepsis and other systemic inflammatory diseases.

## Materials and Methods

### Bacteria and media

L. *plantarum* TW1-1, *Pediococcus acidilactici* XS40, L. *plantarum* DS45, L. *paracasei* LZU-D2, L. *delbruckii,* L. *casei* 18-10, *Streptococcus thermophilus* 19 were provided by Dr. Xusheng Guo (Lanzhou University, Lanzhou, China) which were isolated from yogurt. Bacterial strains were cultured in De Man, Rogosa, and Sharpe (MRS; Beijing Solarbio Science & Technology, Beijing, China) growth medium with exception of 19 and XS40, which were cultured in M17 growth medium (MRS; Beijing Solarbio Science & Technology, Beijing, China) supplemented with 1% lactose and MRS medium supplemented with 0.5% glucose, respectively. MRS and M17 agar medium (Beijing Solarbio Science & Technology, Beijing, China) were used to determine the CFU of the assayed probiotic strains.

### In vitro evaluation of inflammatory factors induced by probiotics

The commercial immortal mouse macrophage cell line RAW264.7 was obtained from the American Type Culture Collection and was grown in Dulbecco’s Modified Eagle’s Medium (DMEM; Gibco, Gaithersburg, MD) supplemented with 10% heat-inactivated fetal bovine serum (FBS) under a humidified 10% CO_2_ atmosphere at 37°C. In order to investigate the influence of probiotics, the cells were cultured in 12-well culture plates at 1×10^6^ cells/well. The bacterial strains were grown in MRS or M17 medium overnight (16 h), after which the cultures were diluted to an optical density (OD) of 0.3, washed with phosphate-buffered saline (PBS; pH 7.4), resuspended in PBS, and were used to infect the RAW264.7 cells at a multiplicity of infection (MOI) of 1:100 (cells/bacteria). The plates were incubated for 6 h at 37°C under a 10% CO2 atmosphere and samples were collected to assess the levels of inflammatory factors by qRT-PCR. PBS without bacteria was used as negative control.

### Animals and sepsis model

The 7-14-week-old BALB/c (H-2D^d^) mice (average weight 20g) used in this study were originally purchased from the Experimental Animal Center of The Fourth Military Medical University and were bred in our facility under specific-pathogen-free conditions. All animals were maintained under a 12 h light/dark cycle. In order to investigate the effect of Lps on the survival rate, mice were administered different doses of lipopolysaccharide (Lps) by intraperitoneal injection. To investigate the influence of probiotics on sepsis, mice were administered 1mg/kg lipopolysaccharide (Lps) by intraperitoneal injection, with a second dose administered 4 days after the first injection. The details of the experimental design are shown in Table 1. The names of the experimental groups were renamed because of sequencing requirements as follows: Lps7 denotes Lps+ S. *thermophilus* 19, 7 denotes S. *thermophilus* 19. All procedures and protocols used in this study conform to the institutional guidelines and were approved by the Ethics Committee of the Fourth Military Medical University.

**Table 1.**
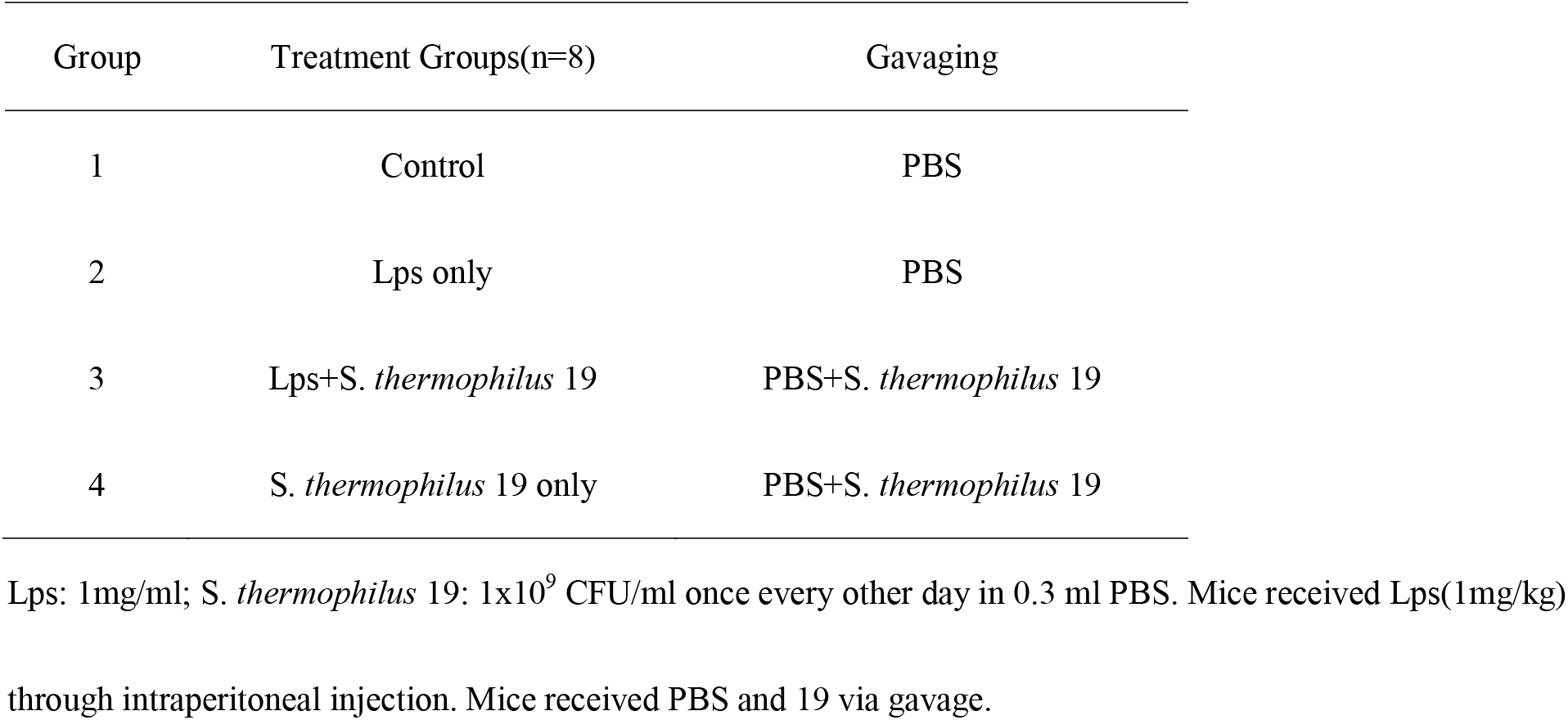
Experimental design

### Weight, water and food intake measurements and sampling

Body weight, water and food intake, and stool appearance were documented for all groups of mice every other day throughout the experiment. After 1 week, livers, kidneys, lungs and small intestines were collected from each mouse and were divided into triplicate samples, with one stored in liquid nitrogen, a second stored in RNAiso Plus for RNA extraction, and the third was fixed in 4% (w/v) paraformaldehyde at 4°C for later histological analysis.

### Histology of different tissues

After the animals were sacrificed, different tissue samples were collected. After fixation in 4% paraformaldehyde, tissue samples were embedded in paraffin and serially cut into 7-mm thick sections. Tissue slides were stained with hematoxylin and eosin (H&E) for histological analysis.

### Microbial DNA extraction and Illumina MiSeq sequencing

Microbial DNA was extracted from the samples using an E.Z.N.A.^®^ Stool DNA Kit (Omega BioTek, Norcross, GA, USA) according to manufacturer’s protocols, and the DNA samples were assessed via PCR with the universal 16S rRNA primers 27F/1492R in our own lab. The DNA concentration and integrity were determined by electrophoresis on 1% agarose gels containing ethidium bromide and spectrophotometrically using an EPOCH instrument (BioTek). After confirmation, the DNA was lyophilized and sent for Illumina MiSeq sequencing and data analysis.

The gut microbiota compositions of mice were assessed via Illumina MiSeq sequencing (Genergy Biotech) targeting the V3-V4 region of the bacterial 16S ribosomal RNA gene using the primers 341F (5’-CCTACGGGNGGCWGCAG-3’) and 785R (5’-GACTACHVGGGTATCTAATCC-3’), with an eight-base barcode sequence unique to each sample. The amplicons were extracted from 2% agarose gels and purified using an AxyPrep DNA Gel Extraction Kit (Axygen Biosciences, Union City, CA, USA) according to the manufacturer’s instructions and were subsequently quantified using a QuantiFluor™-ST instrument (Promega, USA). The purified amplicons were pooled in equimolar ratios and paired-end sequenced (2 × 300) on an Illumina MiSeq platform according to standard protocols. The raw reads were deposited at the NCBI Sequence Read Archive (SRA) database. Operational taxonomic units (OTUs) were clustered with a 97% similarity cutoff using UPARSE (version 7.1 http://drive5.com/uparse/), and chimeric sequences were identified and removed using UCHIME. The taxonomy of each 16S rRNA gene sequence was analyzed using RDP classifier (http://rdp.cme.msu.edu/) against the SILVA (SSU123) 16S rRNA database using a confidence threshold of 70%. The taxonomy of each ITS gene sequence was analyzed using Unite classifier (https://unite.ut.ee/index.php).

### Quantitative RT-PCR for inflammatory factor determination

Total RNA was extracted from different tissues using RNAiso Plus (Takara, Dalian, China) and was subsequently reverse transcribed into cDNA using PrimeScript™ RT Kit (Takara, Dalian, China) according to the manufacturer’s protocol. The expression of inflammatory factor-related genes was analyzed using SYBR^®^ PremixEx Taq™ II and the Bio-Rad CFX system. For real-time PCR, the reaction mixtures contained 1 μL cDNA, 0.4 μL of each primer (10 mmol^-1^), 5 μL of SYBR green PCR Master Mix, and distilled water to a final reaction volume of 10 μL. The Taq DNA polymerase was activated at 95°C for 10 min, followed by 40 cycles of 95°C for 15 s, 60°C for 30 s, and 72°C for 30 s. Quantitative RT-PCR data were normalized to the expression of the housekeeping gene β-actin using the 2^-ΔCt^ method. Primers used in this study are shown in Table 2.

**Table 2.**
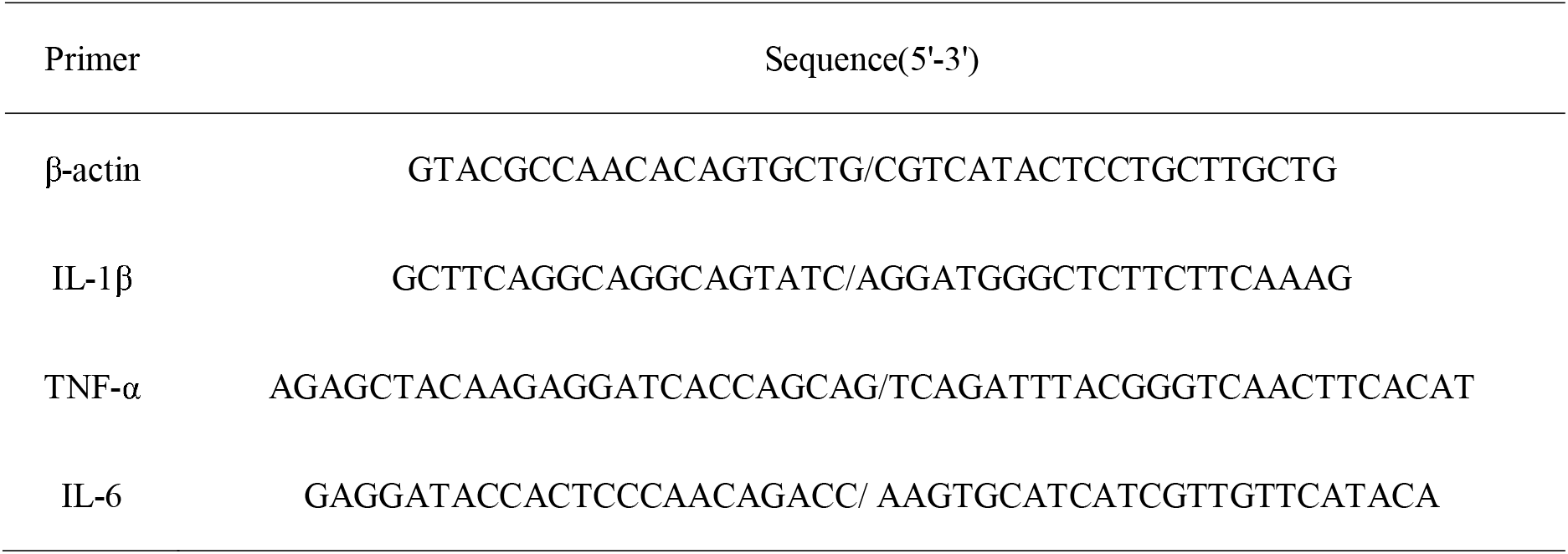
Primers used in this study

### Quantification and statistical analysis

Graphpad Prism was used for graphical presentation and statistical analyses. Differences were considered statistically significant at p<0.05, and data are presented as the means ± SEM. The number of biological replicates (n) and the number of independent experiments are indicated in the figure legends. The Kruskal-Wallis/Wilcoxon rank-sum test was used to analyze the gut microbiota composition data for all the groups.

## Acknowledgements

We thank Dr. Xusheng Guo (Lanzhou University) for his assistance for supply our strains and Dr.Guodong Yang (Fourth Military Medical University) for his advice on data and statistical analysis. This work was supported by grants from Natural Science Foundation of China (81530064, 81801913, 81601680 and 81701902).

## Author contributions

G.W. and D.H. designed and supervised the study. F.H., Y.Z. and X.Y. performed experiments and wrote the manuscript. S.H., Z.F., X.L. and W.C. conducted the animal trial and samples collection. D.X., W.Z. and J.L. helped with animal experiments and provided critical experimental materials and X.Y. conducted physiological data analysis. G.W., D.H. and Y.Z. analysed the data and edited the manuscript.

## Competing interests

The authors declare that they have no conflict of interests

---

**Supplementary Figure1.** The effect of S. *thermophilus* 19 on cell viability was detected by CCK8 assay after co-culture 6hours. Error bars represents SEM.

**Supplementary Figure2.** The influence of S. *thermophilus* 19 and Lps on body weight, toal rat chow and drinking water intake. (A) Body weight change and relative weight change (n=8/group). (B) Total rat chow intake and drinking water.

**Supplementary Figure3.** The details of the change of gut microbiota at phylum (A) and genus (B) level in different groups. Data with significant changes were showed in the figure (P<0.05).

**Supplementary Figure4.** Specific OTUS existed in different groups.

**Supplementary Figure5.** The gut microbiota composition between control group and co-treatment (19 and Lps) group (n=8). (A) (B) Fecal microbiota alfa diversity. (C) PLS_DA plot of fecal microbiota of Lps-treated mice with S. *thermophilus* 19 treatment and control group. (D) The change of gut microbiota at phylum level.

## References

1. Bai X, He T, Liu Y, Zhang J, Li X, Shi J, Wang K, Han F, Zhang W, Zhang Y, Cai W, Hu D. 2018. Acetylation-dependent regulation of notch signaling in macrophages by sirt1 affects sepsis development. Front Immunol 9:762.

2. Manning J. 2018. Sepsis in the burn patient. Crit Care Nurs Clin North Am 30:423–430.

3. Glenwright AJ, Pothula KR, Bhamidimarri SP, Chorev DS, Basle A, Firbank SJ, Zheng H, Robinson CV, Winterhalter M, Kleinekathofer U, Bolam DN, van den Berg B. 2017. Structural basis for nutrient acquisition by dominant members of the human gut microbiota. Nature 541:407–411.

4. Hand TW. 2016. The role of the microbiota in shaping infectious immunity. Trends Immunol 37:647–658.

5. Maynard CL, Elson CO, Hatton RD, Weaver CT. 2012. Reciprocal interactions of the intestinal microbiota and immune system. Nature 489:231–41.

6. Clemente JC, Manasson J, Scher JU. 2018. The role of the gut microbiome in systemic inflammatory disease. BMJ 360:j5145.

7. Carrico CJ, Meakins JL, Marshall JC, Fry D, Maier RV 1986. Multiple-organ-failure syndrome. Arch Surg 121:196–208.

8. Isolauri E, Kirjavainen PV, Salminen S. 2002. Probiotics: a role in the treatment of intestinal infection and inflammation? Gut 50 Suppl 3:III54–III59.

9. Mombaerts P, Mizoguchi E, Grusby MJ, Glimcher LH, Bhan AK, Tonegawa S. 1993. Spontaneous development of inflammatory bowel disease in T cell receptor mutant mice. Cell 75:274–82.

10. Doron S, Snydman DR. 2015. Risk and safety of probiotics. Clin Infect Dis 60 Suppl 2:S129–S134.

11. Wu G, Xiao X, Feng P, Xie F, Yu Z, Yuan W, Liu P, Li X. 2017. Gut remediation: a potential approach to reducing chromium accumulation using *Lactobacillus plantarum* TW1-1. Sci Rep 7:15000.

12. Hwang IY, Koh E, Wong A, March JC, Bentley WE, Lee YS, Chang MW. 2017. Engineered probiotic *Escherichia coli* can eliminate and prevent *Pseudomonas aeruginosa* gut infection in animal models. Nat Commun 8:15028.

13. Correa NB, Peret Filho LA, Penna FJ, Lima FM, Nicoli JR. 2005. A randomized formula controlled trial of *Bifidobacterium lactis* and *Streptococcus thermophilus* for prevention of antibiotic-associated diarrhea in infants. J Clin Gastroenterol 39:385–9.

14. Saavedra JM, Bauman NA, Oung I, Perman JA, Yolken RH. 1994. Feeding of Bifidobacterium bifidum and Streptococcus thermophilus to infants in hospital for prevention of diarrhoea and shedding of rotavirus. Lancet 344:1046–9.

15. Mater DD, Bretigny L, Firmesse O, Flores MJ, Mogenet A, Bresson JL, Corthier G. 2005. *Streptococcus thermophilus* and *Lactobacillus delbrueckii* subsp. bulgaricus survive gastrointestinal transit of healthy volunteers consuming yogurt. FEMS Microbiol Lett 250:185–7.

16. Khosravi A, Yanez A, Price JG, Chow A, Merad M, Goodridge HS, Mazmanian SK. 2014. Gut microbiota promote hematopoiesis to control bacterial infection. Cell Host Microbe 15:374–81.

17. Schuijt TJ, Lankelma JM, Scicluna BP, de Sousa e Melo F, Roelofs JJ, de Boer JD, Hoogendijk AJ, de Beer R, de Vos A, Belzer C, de Vos WM, van der Poll T, Wiersinga WJ. 2016. The gut microbiota plays a protective role in the host defence against pneumococcal pneumonia. Gut 65:575–83.

18. Ojima M, Motooka D, Shimizu K, Gotoh K, Shintani A, Yoshiya K, Nakamura S, Ogura H, Iida T, Shimazu T. 2016. Metagenomic analysis reveals dynamic changes of whole gut microbiota in the acute phase of intensive care unit patients. Dig Dis Sci 61:1628–34.

19. Zaborin A, Smith D, Garfield K, Quensen J, Shakhsheer B, Kade M, Tirrell M, Tiedje J, Gilbert JA, Zaborina O, Alverdy JC. 2014. Membership and behavior of ultra-low-diversity pathogen communities present in the gut of humans during prolonged critical illness. MBio 5:e01361–e14.

20. Pantoflickova D, Corthesy-Theulaz I, Dorta G, Stolte M, Isler P, Rochat F, Enslen M, Blum AL. 2003. Favourable effect of regular intake of fermented milk containing *Lactobacillus johnsonii* on *Helicobacter pylori* associated gastritis. Aliment Pharmacol Ther 18:805–13.

21. Xin J, Zeng D, Wang H, Ni X, Yi D, Pan K, Jing B. 2014. Preventing non-alcoholic fatty liver disease through *Lactobacillus johnsonii* BS15 by attenuating inflammation and mitochondrial injury and improving gut environment in obese mice. Appl Microbiol Biotechnol 98:6817–29.

22. Chiva M, Soriano G, Rochat I, Peralta C, Rochat F, Llovet T, Mirelis B, Schiffrin EJ, Guarner C, Balanzo J. 2002. Effect of *Lactobacillus johnsonii* La1 and antioxidants on intestinal flora and bacterial translocation in rats with experimental cirrhosis. J Hepatol 37:456–62.

23. Borges NA, Stenvinkel P, Bergman P, Qureshi AR, Lindholm B, Moraes C, Stockler-Pinto MB, Mafra D. 2018. Effects of probiotic supplementation on trimethylamine-N-oxide plasma levels in hemodialysis patients: a pilot study. doi: 10.1007/s12602-018-9411-1. Probiotics Antimicrob Proteins:1–7.

24. Sonnenburg JL, Backhed F. 2016. Diet-microbiota interactions as moderators of human metabolism. Nature 535:56–64.

25. Wang J, Tang H, Zhang C, Zhao Y, Derrien M, Rocher E, van-Hylckama Vlieg JE, Strissel K, Zhao L, Obin M, Shen J. 2015. Modulation of gut microbiota during probiotic-mediated attenuation of metabolic syndrome in high fat diet-fed mice. ISME J 9:1–15.

