## Supplementary figures and images for "Remodeling gut microbiota by *Streptococcus thermophilus* 19 attenuates inflammation in septic mice"

### Supplementary Figure1

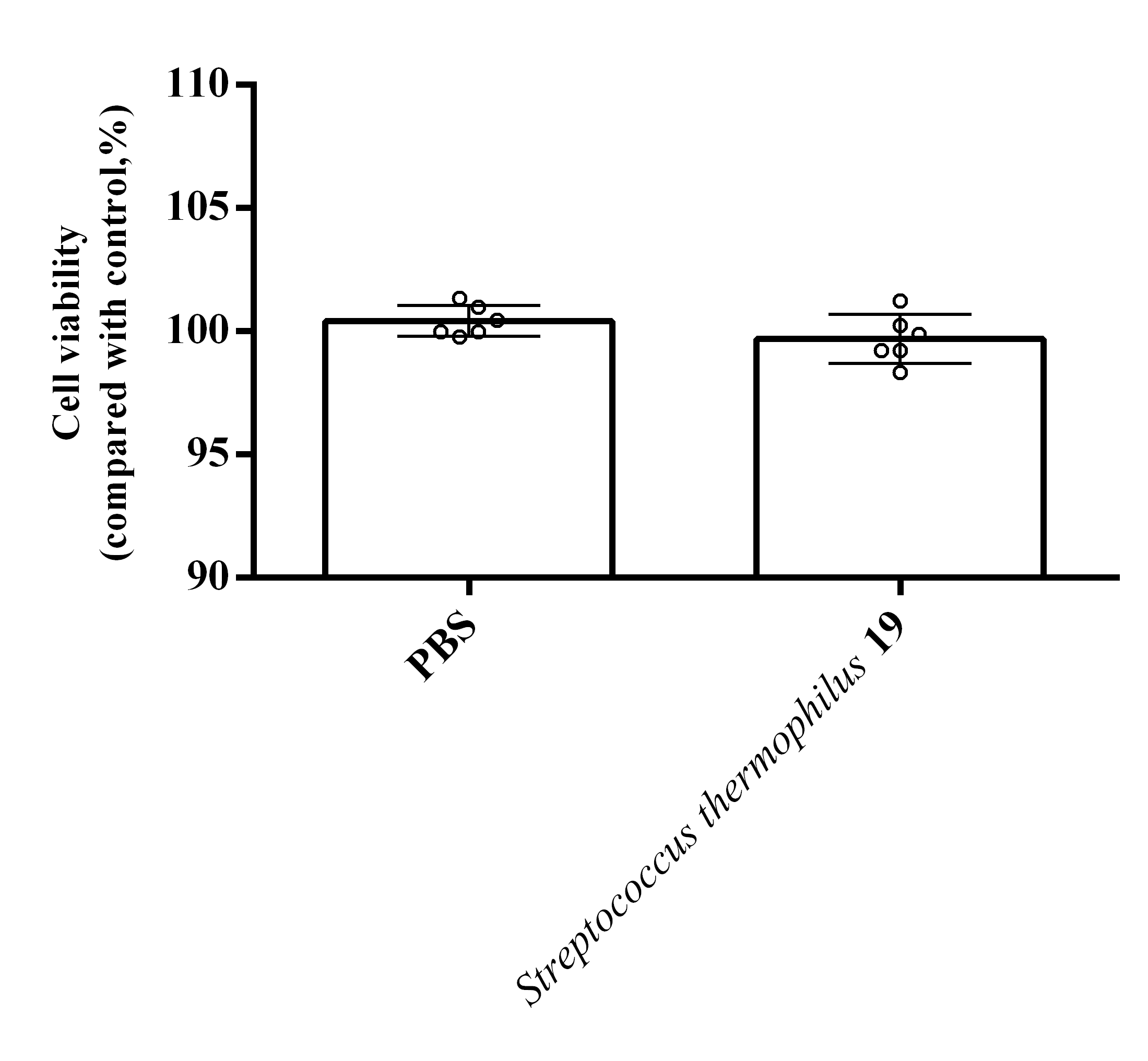

### Supplementary Figure2

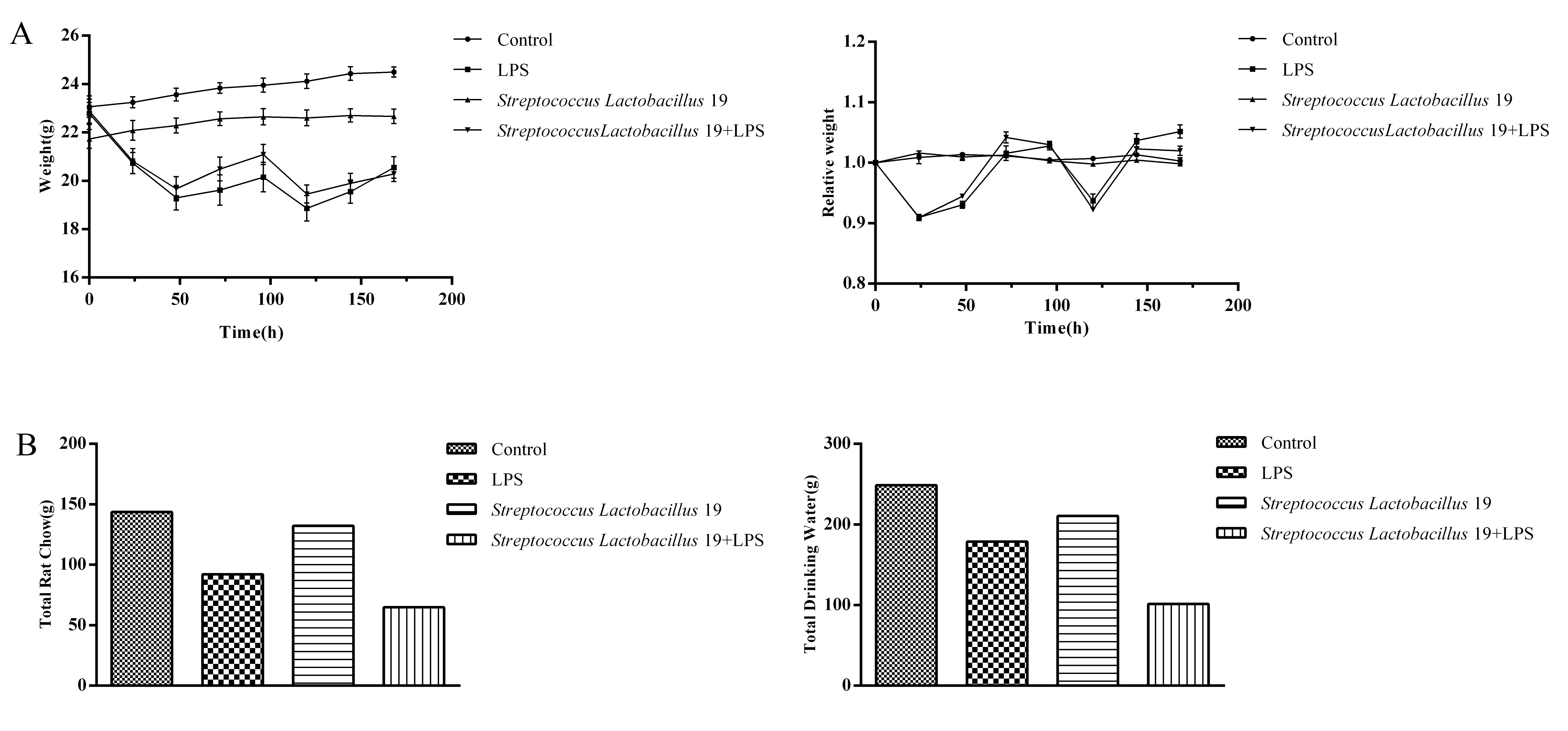

### Supplementary Figure3

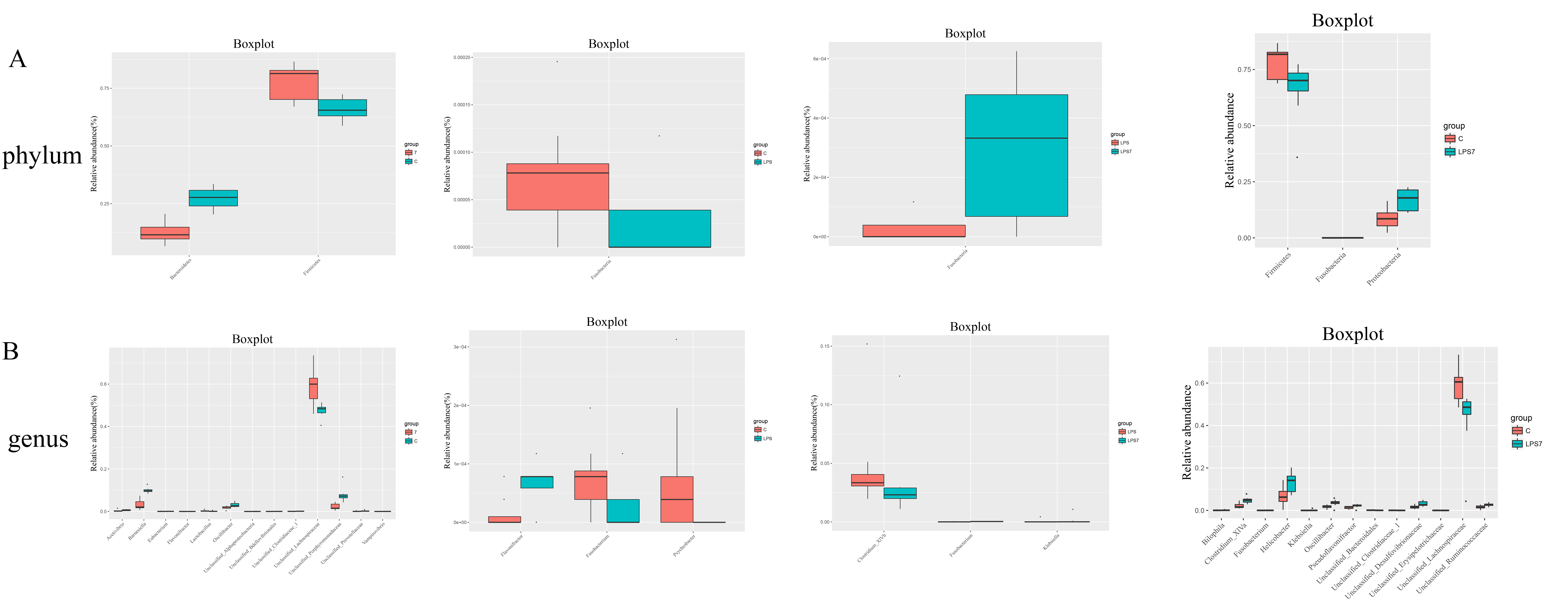

### Supplementary Figure4

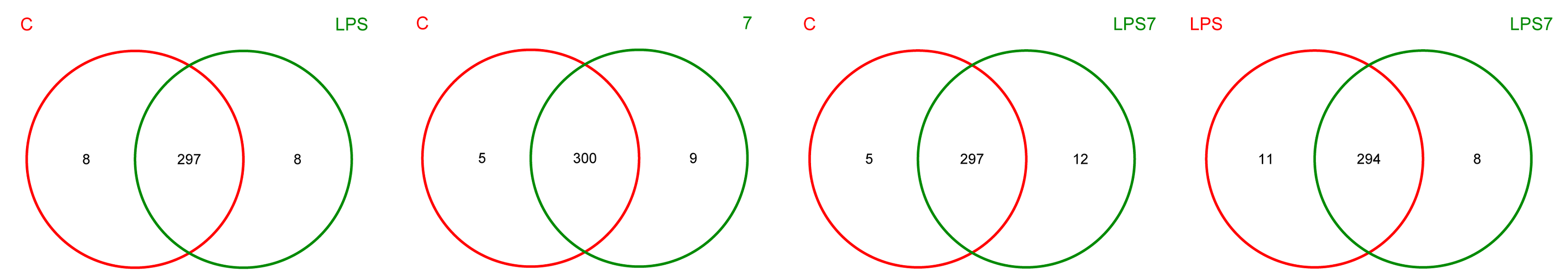

### Supplementary Figure5

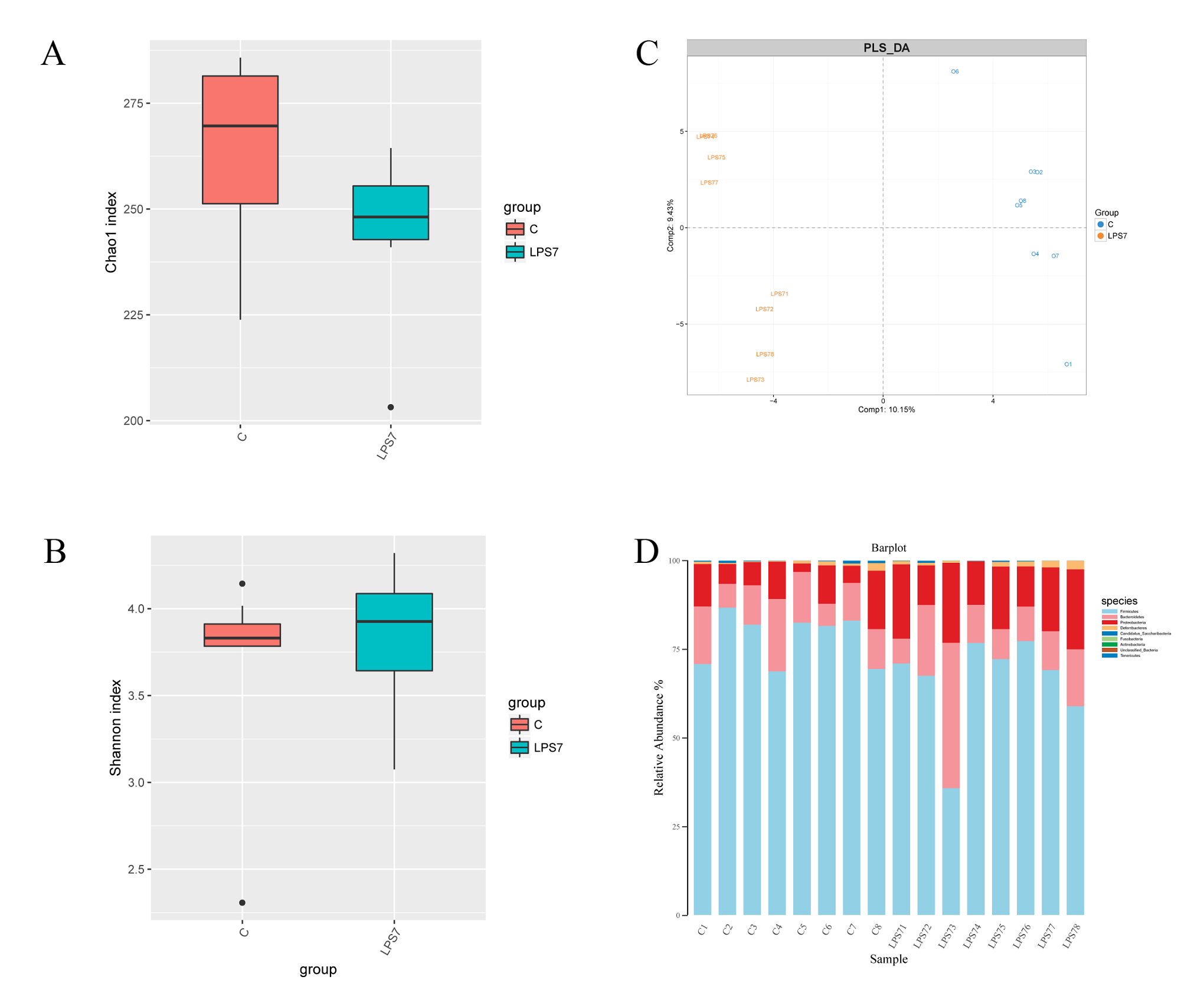
